# Complementation between pathological prion protein subassemblies to cross existing species barriers

**DOI:** 10.1101/861278

**Authors:** Angélique Igel-Egalon, Florent Laferrière, Philippe Tixador, Mohammed Moudjou, Laetitia Herzog, Fabienne Reine, Juan Maria Torres, Hubert Laude, Human Rezaei, Vincent Béringue

**Affiliations:** VIM, INRA, Université Paris-Saclay, 78350, Jouy-en-Josas, France; Centro de Investigación en Sanidad Animal (CISA-INIA), Valdeolmos, Madrid, Spain

**Keywords:** Prion, species barrier, transgenic mice, quasi-species, sedimentation velocity, assemblies

## Abstract

**Background:** prion replication results from the autocatalytic templated assisted conversion of the host-encoded prion protein PrP_C_ into misfolded, polydisperse PrPSc conformers. Structurally distinct PrP_Sc_ conformers can give rise to multiple prion strains. Within and between prion strains, the biological activity (replicative efficacy and specific infectivity) of PrP_Sc_ assemblies is size-dependent and thus reflects an intrinsic structural heterogeneity. The contribution of such PrP_Sc_ heterogeneity across species prion adaptation, - which is believed to be based on fit-adjustment between PrP_Sc_ template(s) and host PrP_C_ -, has not been explored.

**Methods:** to define the structural-to-fitness PrP_Sc_ landscape, we measured the relative capacity of size-fractionated PrP_Sc_ assemblies from different prion strains to cross mounting species barriers in transgenic mice expressing foreign PrP_c_.

**Results:** in the absence of a transmission barrier, the relative efficacy of the isolated PrP_Sc_ assemblies to induce the disease is superimposable to the efficacy observed in the homotypic context. However, in the presence of a transmission barrier, size fractionation overtly delays and even abrogates prion pathogenesis in both neural and extraneural, prion-permissive tissues, for reason independent of the infectivity load of the isolated assemblies. This suggests that a synergy between structurally distinct PrP_Sc_ assemblies in the inoculum is requested for crossing the species barrier. We further strengthen this hypothesis by showing that altering, by serial dilution, PrP_Sc_ assemblies content of unfractionated inocula reduce their specific infectivity in an aberrant manner, solely in the presence of a transmission barrier.

**Conclusions:** our data support a mechanism whereby overcoming prion species barrier requires complementation between structurally distinct PrP_Sc_ assemblies. This work provides key insight into the “quasi-species” concept applied to prions, which would not necessarily rely on prion sub-strains as constituent but on structural PrP_Sc_ heterogeneity within prion population.

## Background

Mammalian prions are proteinaceous pathogens formed from misfolded assemblies (PrP_Sc_) of the host-encoded prion protein PrP_C_. Prions self-replicate by templating the conversion and polymerization of PrP_C_ [1]. Prions cause inexorably fatal neurodegenerative diseases such as human Creutzfeldt-Jakob disease (CJD), sheep scrapie, bovine spongiform encephalopathy (BSE) and chronic wasting disease of cervids [2].

The prion strain phenomenon which is due to a structural polymorphism of PrP_Sc_ assemblies is defined by the physiopathological and biochemical characteristics of prion disease within these host species and in experimental models [3–10]. The strain-specified physiopathological differences include duration of disease in the challenged or affected species, vacuolation distribution and pattern of PrP_Sc_ deposition in the brain, tropism for the lymphoid tissue and biochemical properties of PrP_Sc_, including resistance to denaturation and to proteases [3]. How PrP_Sc_ structural polymorphism causes such distinct phenotypes remains poorly understood. A broad panel of experimental observations, including by size fractionation supports the existence of structurally different PrP_Sc_ subsets within a given prion strain [11–19]. This intra-strain structural heterogeneity results from the intrinsic and deterministic properties of the prion replication process to generate structurally diverse PrP_Sc_ subsets [20]. Structural diversity can be observed at different levels. Size-fractionation studies by sedimentation velocity (SV) indicate that variability in PrP_Sc_ quaternary structure is strain-specific [15,18,17,21,22]. Within a given prion strain, PrP_Sc_ assemblies with differing quaternary structure exhibit markedly different templating and biological activities, which are a hallmark of the existence of structurally distinct PrP_Sc_ subpopulations, as previously discussed [23]. The biological and biochemical consequences of PrP_Sc_ structural heterogeneity during prion replication and propagation are poorly understood.

Maybe one of the most intriguing and unpredictable change in prion replicative environment is the interspecies transmission. In certain host/strain combinations, prions will propagate readily as in an intraspecies transmission. In other host/strain combinations, only a fraction of the exposed animals will develop the disease with variable incubation periods, reflecting a ‘species’ or ‘transmission’ barrier. Attaining full attack rate and minimal incubation periods will require iterative transmissions. At that stage, prions are considered to be adapted to the new host [3]. Two main theoretical models have been developed to describe the transmission barrier at the molecular level. Both propose that the species barrier is governed by a misfit between PrP_Sc_ contained in the infecting prion and host PrP_C_ folding landscape [5,24]. Schematically, the first one called ‘deformed templating’ consider that PrP_C_ will progressively adopt, -due to its specific conformational dynamic-, the “quasi-right” conformation to be selected during the templating process [24]. As a consequence, a switch to a new strain structural determinant can occur, which combines structural information from the inoculated PrP_Sc_ assemblies with the folding landscape of the new host PrP_C_ [25–29]. A major limitation of this model resides in the assumption that all PrP_Sc_ assemblies are structurally equivalent, despite experimental counter-evidence. The second model called ‘conformational selection’ considers the existence of structurally distinct PrP_Sc_ subsets within a strain or isolate, the species adaptation resulting from the selection of the best replicator [5]. The two models are not mutually exclusive. They at least have the merit to highlight the role of structural diversity of PrP_C_ and/or PrP_Sc_ in prion adaptation and evolution.

Here, we aim at defining the contribution of PrP_Sc_ (quaternary) structure polydispersity to across species prion fitness. We compare the relative capacity of size-fractionated PrP_Sc_ assemblies to propagate in transgenic mice expressing homotypic versus heterotypic PrP_C_, used as proxy of mounting species barriers (one with prion ‘mutation’). We show that fractionating PrP_Sc_ assemblies strengthens existing species barrier, to a degree of magnitude independent of their infectivity load. Coexistence of heterogeneous PrP_Sc_ quaternary structure in the inoculum may thus be requested to cross species barriers. We strengthen this hypothesis by showing that altering, by serial dilution, PrP_Sc_ assemblies content overtly reduces the specific infectivity from unfractionated inocula during cross-species transmission events. Our data support a mechanism whereby overcoming prion species barrier requires complementation between structurally distinct PrP_Sc_ subsets within the inoculum, refining the current prion adaptation models.

## Methods

### Transgenic mouse lines

The transgenic mouse lines expressing ovine, hamster and bovine PrP have been described previously [30–33]. The bovine PrP tg540 [30] and tg110 [31] lines showed equivalent susceptibilities to classical BSE prions [30].

### Prion sources

LA19K, LA21K *fast* and 127S scrapie prions are cloned prion strains. They have been obtained by serial transmission and subsequent biological cloning by limiting dilutions of classical field scrapie isolates to *tg338* transgenic mice expressing the VRQ allele of ovine PrP [32,18]. Pooled *tg338* mouse brain homogenates (20% wt/vol. in 5% glucose) were used in centrifugation analyses, as indicated.

The transmission properties of the L-type BSE isolate (designated BASE by the authors [34]) in tgBov mice (tg540 line) and tgOv mice have been previously described [26]. The same brain material from the same isolate was transmitted to tgBov line (tg110) line by intracerebral route by using 20 µl of a 10% (wt/vol. in 5% glucose) brain homogenate.

### Fractionation by sedimentation velocity

The sedimentation velocity procedure has been previously and comprehensively described [15,18,20,13]. Briefly, detergent-solubilized brain homogenates were loaded atop a continuous 10–25% iodixanol gradient (Optiprep, Axys-shield). The gradients were centrifuged at 285 000 g for 45 min in a swinging-bucket SW-55 rotor using an Optima LE-80K ultracentrifuge (Beckman Coulter). Gradients were then manually segregated into 30 equal fractions of 165µl from the bottom using a peristaltic pump. Fractions were aliquoted for immunoblot and bioassays. Gradient linearity was verified by refractometry. To avoid any cross-contamination, each piece of equipment was thoroughly decontaminated with 5 M NaOH followed by several rinses in deionised water after each gradient collection.

### Bioassays

Fractions were diluted extemporarily in 5% glucose (1:5) in a class II microbiological cabinet according to a strict protocol to avoid any cross-contamination. Individually identified 6- to 10-week-old mice were inoculated intracerebrally with 20 µl of the solution, using a 27-gauge disposable syringe needle inserted into the right parietal lobe. At terminal stage of disease or at end-life, mice were euthanized and analyzed for proteinase K (PK) resistant PrP_Sc_ (PrP_res_) content in brains and spleens tissues (as indicated), using the Bio-Rad TsSeE detection kit [26] before immunoblotting, as described below.

### Endpoint titration

For titration purposes, groups of indicator mice were inoculated intracerebrally (20 µl) with serial ten-fold dilutions of the indicated brain homogenates. Animals inoculated with the initial dose at 10% (w/v) solution were assigned an infectious dose of 0. The mice were monitored daily, euthanized at terminal stage and analyzed as above for PrP_res_ content. The survival times of tgBov or tgHa reporter mice was measured for each tenfold dilution tested and the relative infectious dose / survival time relationship was reported, when available. It allows translating survival times of the inoculated fractions in infectious dose (i.e., equivalent to that found in brain homogenate dilutions) as estimate of infectivity.

### Immunoblots

Aliquots of the collected fractions were treated with a final concentration of 50 µg/ml PK (1 hour, 37°C). Samples were then mixed in Laemmli buffer and denatured at 100°C for 5 min. The samples (15 µl) were run on 12% Bis-Tris Criterion gels (Bio-Rad, Marne la Vallée, France) and analyzed by immunoblots, using the Sha31 anti-PrP antibody (human PrP epitope at residues 145 to 152 [35]). Immunoreactivity was visualized by chemiluminescence (GE Healthcare). The amount of PrP present per fraction and the PrP_Sc_ glycoforms ratios were determined with the GeneTools software after acquisition of chemiluminescent signals with a GeneGnome digital imager (Syngene, Frederick, Maryland, United States).

### Histoblots

For histoblotting procedure, brains were rapidly removed from euthanized mice and frozen on dry ice. Cryosections were cut at 8–10 μm, transferred onto Superfrost slides and kept at −20°C until use. Histoblot analyses were performed on 3 brains per experiment, using the 12F10 anti-PrP antibody (human PrP epitope at residues 145 to 160 [36]).

## Results

### Transgenic modelling of prion species barrier

We used a previously developed SV protocol to separate different populations of PrP_Sc_ assemblies according to their quaternary structure [13,15,18]. To determine the potential role of the SV-isolated PrP_Sc_ subpopulations in prion adaptation and evolution, we included in the present study three well-characterized ovine prion strains termed LA19K, LA21K *fast* and 127S. These cloned strains markedly differ according to the specific infectivity of their respective PrP_Sc_ subpopulations in the homotypic PrP transmission conditions. For LA21K *fast* and 127S strains, a discrete population of small oligomers (<pentamers) exhibit the highest specific infectivity values. For LA19K, the highest specific infectivity values are associated with larger-size oligomers (>40 PrP-mers) [23]. We also included atypical L-BSE prions in the study, because of their mutability on cross-species transmission [37].

We transmitted SV fractionated-PrP_Sc_ assemblies from the aforementioned strains to transgenic mouse models expressing heterotypic PrP. The models were chosen according to the transmission barrier observed with unfractionated material, as examined by gold-standard criteria [38–40], including disease attack rate on primary passage, reduction of incubation durations (ID) on serial passaging, and establishment of prion strain properties. The same analyses were performed on back-passage to mice expressing the parental host PrP_C_. For the sake of clarity, these data are comprehensively summarized as Additional file 1, supplementary text and supplementary Fig. 1-4. However, to provide a straightforward estimator of the stringency of the species barrier, we used the reduction factor between the mean ID on first to second passage in the heterotypic context and on retrotransmission [40]. These data are shown as Fig. 1a. A magnitude of 1 signifies prion straight adaptation. As model of prion transmission without species barrier, LA19K scrapie prions from ovine PrP (tgOv) mice was passed to bovine PrP mice (tgBov). As model of prion transmission with species barrier and ‘mutation’, we passed L-BSE prions onto tgOv mice. As models of prion transmission with strong species barrier, we passed LA21K *fast* and 127S scrapie prions onto hamster PrP mice (tgHa).

**Figure 1.**
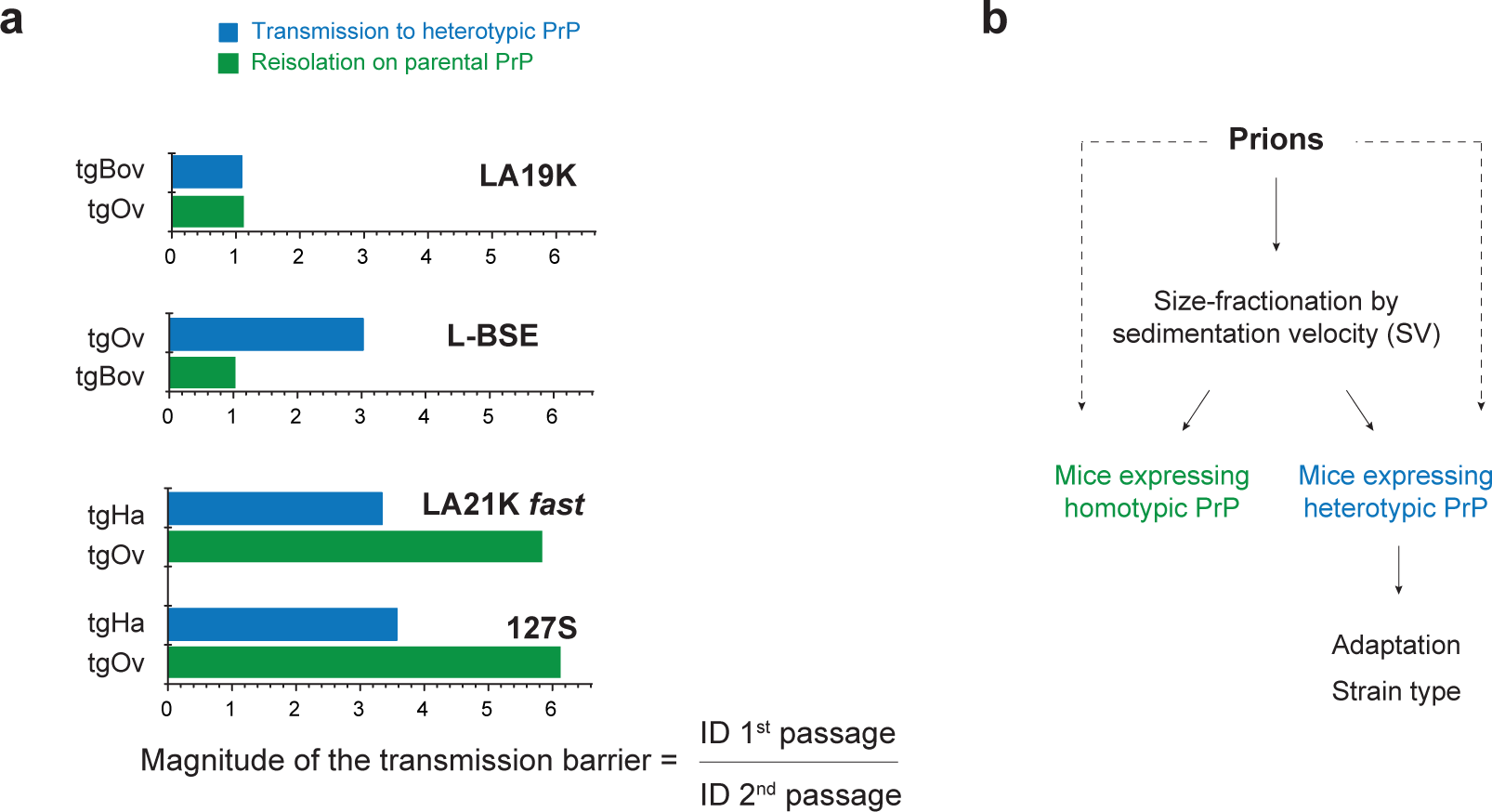
Magnitude of the species barrier on transmission of scrapie prions (LA19K, LA21K *fast*, 127S) and cattle prions (L-BSE) to transgenic mice expressing heterotypic PrP. (**a**) Scrapie LA19K, LA21K *fast*, 127S prions and cattle L-BSE prions were transmitted iteratively to mice expressing bovine PrP (tgBov), hamster PP (tgHa) and ovine PrP (tgOv), respectively, before reisolation in transgenic mice expressing the parental PrP (Additional file 1). The magnitude of the transmission barrier in transgenic mice expressing heterotypic PrP (blue color) and on reisolation in transgenic mice expressing the parental PrP (green color) was calculated as the ratio of the mean incubation durations (ID) on first to second passage in the new host PrP and on reisolation in mice expressing the parental PrP. A magnitude of 1 signifies prion straight adaptation. (**b**) Overview of the bioassays made with PrP_Sc_ assemblies fractionated by sedimentation velocity.

### Impact of size-fractionation on prion pathogenesis in the heterotypic PrP context

Brain homogenates from tgOv mice containing LA19K prions, LA21K *fast* or 127S prions and from cattle containing L-BSE prions were solubilized and fractionated by SV [15,18]. The fractions were then intracerebrally inoculated in the heterotypic PrP_C_ context to determine the specific transmission propensity of each fraction. (Fig. 1b). As controls, some fractions were transmitted in the homotypic PrP context and confirmed our previous results ([18]; *vide infra*). In the absence of any apparent species barrier, as determined for LA19K inoculated to tgBov mice, the infectivity sedimentograms in the homotypic and heterotypic passage conditions, tended to superimpose, in terms of distribution and infectivity values amongst the fractions (Fig. 2a). Each LA19K isolated PrP_Sc_ assembly thus exhibited quasi-equivalent specific infectivity in both transmission contexts. The strain-specified PrP_res_ electrophoretic profile and pattern of cerebral PrP_res_ deposition [33] were conserved amongst the tgBov mice inoculated with the different fractions or unfractionated LA19K brain material (Fig. 2b and additional file 1, supplementary Fig. 5), suggesting phenotypic invariance of the isolated LA19K assemblies on heterotypic transmission.

**Figure 2.**
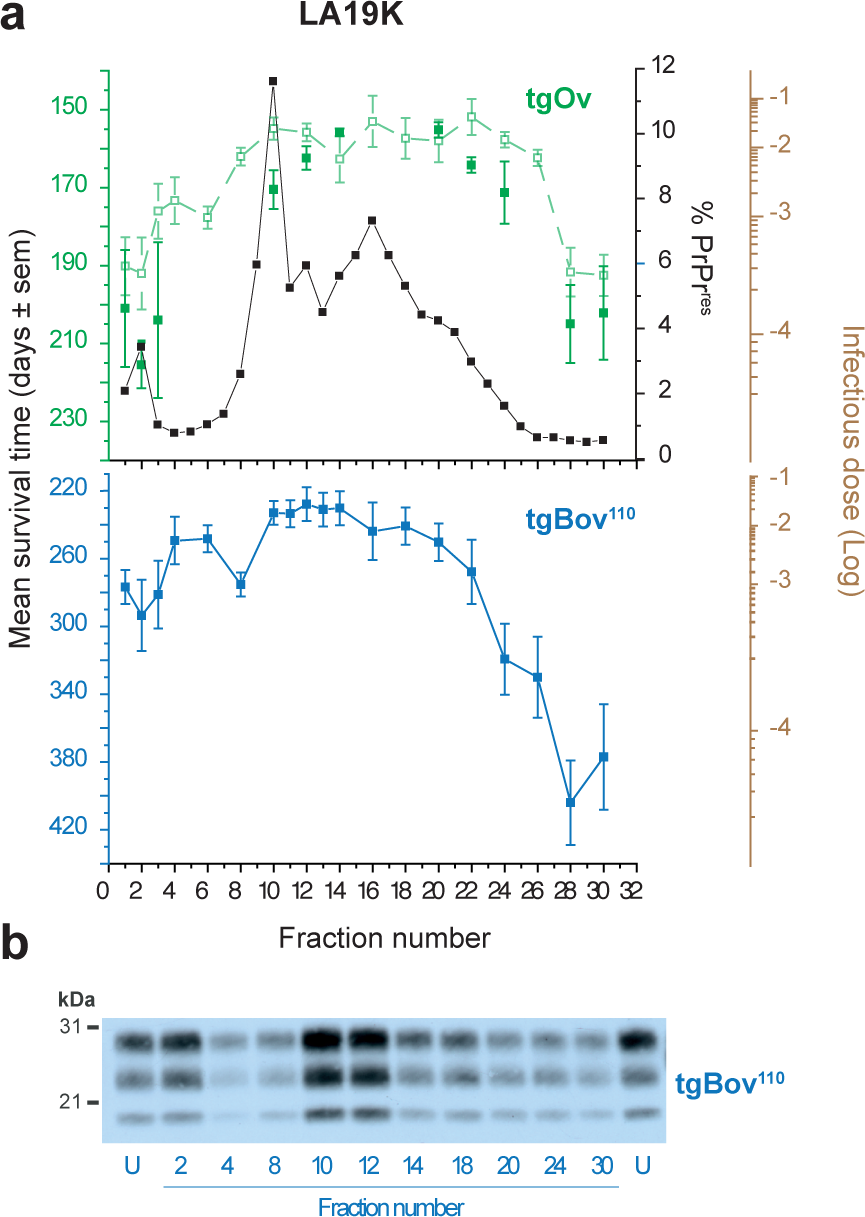
Unaltered capacity of SV-fractionated PrP_Sc_ assemblies to propagate onto a new host PrP sequence in the absence of a transmission barrier. (**a**) SV profiles of PrP_res_ (black line) and infectivity in the homotypic PrP (green line) and heterotypic PrP (blue line) transmission context. Brain homogenates from tgOv mice inoculated with LA19K prions were solubilized before fractionation by SV. The collected fractions were analyzed for PrP_res_ content by immunoblot and for infectivity by an incubation time bioassay in tgOv and tgBov mice (tg110 line). In the homotypic context, plain and dotted symbols/lines refer to this study and to previous reports [18], respectively. The right logarithmic brown scale provides the LA19K-specific reciprocal relation between survival time in tgOv and tgBov mice and infectious dose, as established by limiting dilution titration (as from Table 3 and [18]). Animals inoculated with 10% infectious brain material are assigned an infectious dose of 0. (**b**) PrP_res_ electrophoretic profiles in the brains of tgBov mice inoculated with size-fractionated LA19K-tgOv PrP_Sc_ assemblies. The profile obtained with unfractionated (U) material is shown for comparison.

In the presence of a strong species barrier, as determined for LA21K *fast* and 127S inoculated to tgHa mice, SV-fractionation had a strong negative impact. None of the mice inoculated with the fractions from three independent gradients developed any neurological symptoms up to end-life. Only 3 out of the 221 tested mice (i.e., 1%) accumulated PrP_res_ in the brain. These 3 mice were inoculated with fractions 10 and 13 from one LA21K *fast* gradient (Fig. 2,Table 1). In particular, none of the tgHa mice inoculated with the PrP_Sc_ assemblies with the highest specific infectivity values from fractions 1-2 accumulated any detectable PrP_res_ (n=43, Table 1). These results were confirmed by a second passage with individual or pooled (by 2) tgHa brains, covering the entire LA21K *fast* gradient. All the secondary transmissions with PrP_res_-negative brains were negative (neurological signs and PrP_res_; Table 2), indicating absence of a subclinical disease [41,42] and/or of infectious, PK-sensitive PrP_Sc_ species [16,43] in the non-responder mice. Oppositely, serial transmissions from PrP_res_-positive brains led to isolation of prions with strains properties identical to those obtained on adaptation of unfractionated LA21K *fast* prions to tgHa mice (Table 2, Fig. 2b, Additional file 1, supplementary Fig. 3).

**Table 1.** Survival time of hamster PrP mice inoculated with LA21K *fast* and 127S prions fractionated by sedimentation velocity.

| Fractions | LA21K *fast* |  |  |  | 127S |  |
| --- | --- | --- | --- | --- | --- | --- |
|  | n/n01 | Survival<sub>2</sub><br>(days ± SEM) | n/n01 | Survival<sub>2</sub><br>(days ± SEM) | n/n01 | Survival<sub>2</sub><br>(days ± SEM) |
| **1** | 0/12 | 399-627 | 0/5 | 384-631 | 0/6 | 365-552 |
| **2** | 0/13 | 520-692 | 0/6 | 318-576 | 0/5 | 341-496 |
| **3** | 0/5 | 389-651 | 0/6 | 401-636 | 0/5 | 405-538 |
| **4** | 0/5 | 507-647 | 0/6 | 384-535 | 0/6 | 317-506 |
| **6** | 0/6 | 386-780 |  |  |  |  |
| **8** | 0/6 | 375-713 |  |  |  |  |
| **10** | 2/6 | 353; 386 | 0/6 | 461-531 | 0/6 | 331-510 |
| **11** | 0/5 | 536-937 | 0/6 | 378-517 |  |  |
| **12** | 0/11 | 347-687 | 0/6 | 322-426 | 0/6 | 382-587 |
| **13** | 1/6 | 394 |  |  |  |  |
| **14** | 0/6 | 556-720 | 0/6 | 344-629 | 0/6 | 481-631 |
| **15** | 0/6 | 323-854 |  |  |  |  |
| **16** | 0/6 | 614-710 |  |  |  |  |
| **18** | 0/6 | 325-839 |  |  |  |  |
| **20** | 0/6 | 406-877 |  |  |  |  |
| **22** | 0/6 | 371-751 |  |  |  |  |
| **24** | 0/5 | 543-927 |  |  |  |  |
| **26** | 0/6 | 413-602 |  |  |  |  |
| **28** | 0/6 | 345-749 |  |  |  |  |
| **30** | 0/6 | 383-665 |  |  |  |  |
| **U<sub>3</sub>** | 8/8 | 157 ± 6 |  |  | 8/8 | 168 ± 7 |

n/n0: Number of mice with neurological disease or positive for PrP<sub>res</sub> in the brain by immunoblotting/number of intracerebrally inoculated mice.
<sub>2</sub>For mice negative for PrP<sub>res</sub> accumulation, only the range of survival time is given.
<sub>3</sub>U: unfractionated material (Additional File 1, supplementary Fig. 3).

**Table 2.** Serial passage of SV fractionated LA21K *fast* prions in hamster PrP mice.

| Fractions<br>(mouse<br>no.1) | LA21K *fast* |  |  |  |  |  |
| --- | --- | --- | --- | --- | --- | --- |
|  | 2<sup>nd</sup> passage |  | 3<sup>rd</sup> passage |  | 4<sup>th</sup> passage |  |
|  | n/n<sub>02</sub> | Survival<sub>3</sub> | n/n<sub>02</sub> | Survival<sub>3</sub> | n/n<sub>02</sub> | Survival<sub>3</sub> |
| **1** (no. 2+3) | 0/6 | 382-524 |  |  |  |  |
| **2** (no.4+7) | 0/6 | 301-605 |  |  |  |  |
| **10**<br>(no.1+2) | 6/6 | 54 ± 1 | 6/6 | 50 ± 1 | 6/6 | 48 ± 1 |
| **11**<br>(no.1+4) | 0/6 | 353-544 |  |  |  |  |
| **12**<br>(no.1+2) | 0/6 | 347-687 |  |  |  |  |
| **13** (no.1) | 6/6 | 58 ± 1 | 6/6 | 48 ± 1 |  |  |
| **20**<br>(no.2+5) | 0/6 | 361-518 |  |  |  |  |
| **26**<br>(no.1+3) | 0/6 | 405-587 |  |  |  |  |
| **U<sub>4</sub>** | 6/6 | 48 ± 1 | 6/6 | 47 ± 1 | 6/6 | 46 ± 1 |

<sup>1</sup>mouse identification number inoculated as pools.
<sup>2</sup>n/n<sub>0</sub>: Number of mice with neurological disease or positive for PrP<sub>res</sub> in the brain by immunoblotting/number of intracerebrally inoculated mice (the number of the mouse inoculated is indicated).
<sup>3</sup>For mice negative for PrP<sub>res</sub> accumulation, only the range of survival time is given.
<sup>4</sup>U: unfractionated material (Additional File 1, supplementary Fig. 3).

To discard the possibility that the loss of tgHa-transmissibility of the isolated PrP_Sc_ was due to an insufficient infectivity load, we estimated the infectivity levels of the top and middle fractions of one LA21K *fast* gradient. Fractions 2 and 12 induced disease in tgOv mice in 68 ± 3 days (5/5) and 82 ± 1 days (5/5), respectively (Fig. 3a), meaning, according to LA21K *fast* dose/response curve (Fig. 3a, Table 3), that their infectivity levels were equivalent to 10_-1.2_ and 10_-3.2_ dilutions of LA21K *fast* brain material, respectively. These values were fully consistent with the mean (±SEM) relative infectious dose contained in LA21K *fast* most infectious upper fractions and middle fractions established from seven independent bioassays (top fractions: mean: 10_-1.36_ (10_-1.08_ to 10_-1.64_); middle fractions: between 10_-2.9_ and 10_-4.65_ depending on the middle fractions). To estimate the infectivity load of these factions in the heterotypic PrP context, we established a dose/response curve of LA21K *fast* in tgHa mice by transmitting by intracerebral route serial tenfold dilutions of a LA21K *fast* tgOv-brain to reporter tgHa mice. The limiting dilution value of LA21K *fast* prions in tgHa mice was surprisingly low (*vide infra*), establishing at 10_-2_ (Table 3). Nevertheless, this value was below the infectivity value of the top fractions of the gradient. Those were thus sufficiently infectious *per se* to induce or transmit disease, at least partly in tgHa mice. This was not observed in 3 independent experiments with a sufficient number of inoculated mice. Almost counterintuitively, the sole fractions eliciting asymptomatic replication in a very low number of mice had infectivity values below the limiting dilution. This suggests a stochastic transmission process and lends support to the view that infection is possible at doses below the limiting dilution [44].

**Figure 3.**
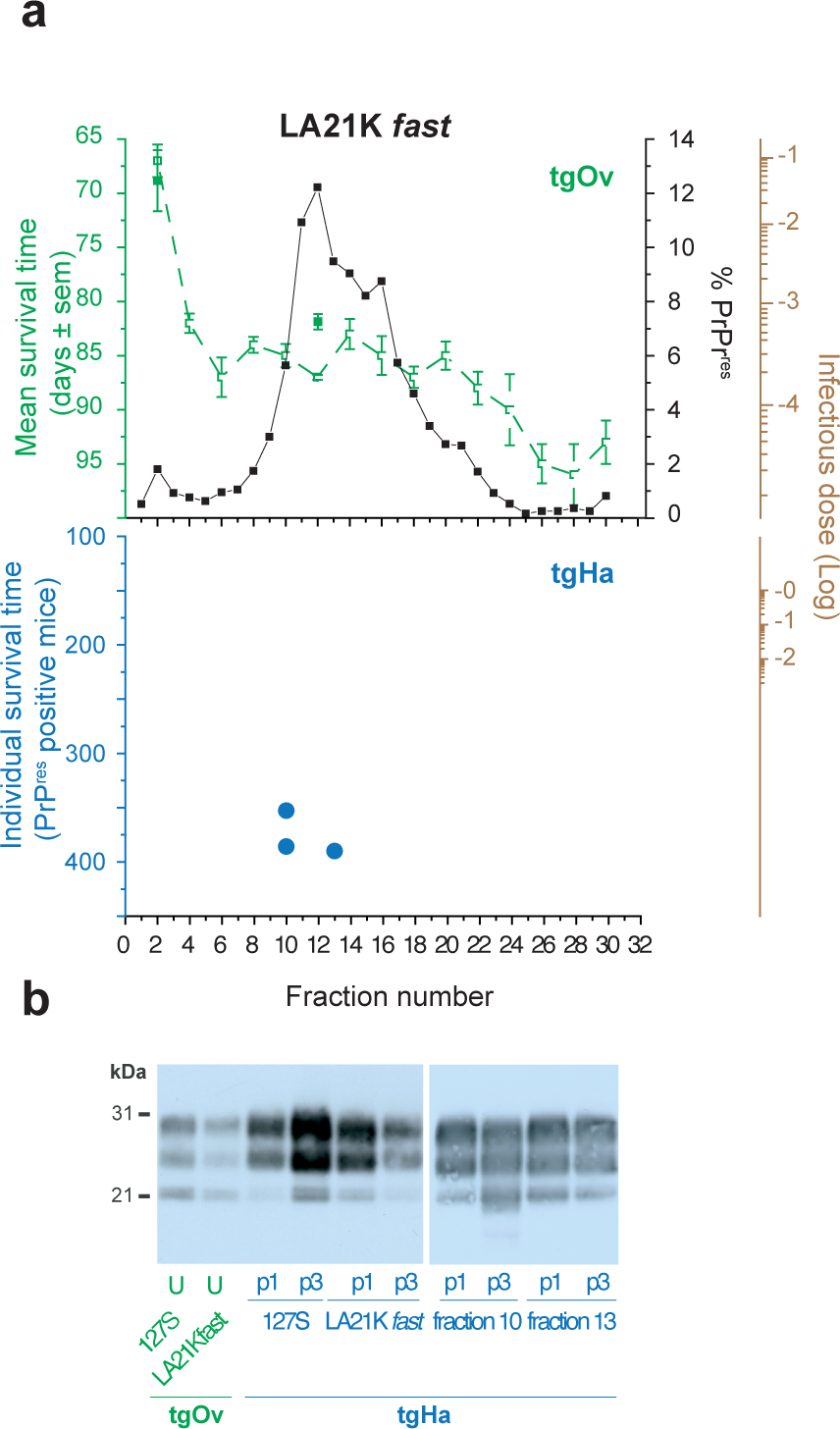
Altered capacity of size-fractionated PrP_Sc_ assemblies to propagate onto a new host PrP sequence in the presence of a ‘strong’ transmission barrier. (**a**) SV profiles of PrP_res_ (black line) and infectivity in the homotypic PrP (green line) and heterotypic PrP (blue dots) transmission context. Brain homogenates from tgOv mice inoculated with LA21K *fast* prions were solubilized before fractionation by SV. The collected fractions were analyzed for PrP_res_ content by immunoblot and for infectivity by an incubation time bioassay in tgOv and tgHa mice. In the homotypic context, plain and dotted symbols/lines refer to this study and to previous reports [18], respectively. Because of the reduced penetrance of the disease in tgHa mice, the mean individual ID of the PrP_res_ positive mice are shown as blue dots. The right logarithmic brown scale provides the LA21K *fast*-specific reciprocal relation between survival time in tgHa and tgOv mice and infectious dose, as established by limiting dilution titration (as from Table 3 and [18]). Animals inoculated with 10% infectious brain material are assigned an infectious dose of 0. (**b**) PrP_res_ electrophoretic profiles in the brains of tgHa mice inoculated with size-fractionated LA21K *fast* prions at the 1st (p1) and 3rd (p3) passage. The profiles obtained with unfractionated (U) LA21K *fast* and 127S material in tgOv and tgHa mice are shown for comparison.

**Table 3.** Endpoint titration of LA19K and LA21K *fast* scrapie prions in transgenic mice expressing homotypic and heterotypic PrP.

| Dilution<sup>1</sup> | Survival time<sup>2</sup> (n/n<sub>0</sub>)<sup>3</sup> |  |  |  |
| --- | --- | --- | --- | --- |
|  | LA19K |  | LA21K *fast* |  |
|  | Ovine PrP | Bovine PrP<sub>4</sub> | Ovine PrP | Hamster PrP |
| 10<sup>-0</sup> | *128 ± 2 (6/6)* | *196 ± 3 (6/6)* | *60 ± 2 (5/5)* | *153 ± 6 (6/6)* |
| 10<sup>-1</sup> | *151 ± 3 (6/6)* | *213 ± 5 (6/6)* | *64 ± 1 (5/5)* | *181 ± 11 (6/6)* |
| 10<sup>-2</sup> | *167 ± 4 (6/6)* | *240 ± 5 (6/6)* | *71 ± 1 (5/5)* | *192; 230 (2/6)* |
| 10<sup>-3</sup> | *188 ± 4 (6/6)* | *272 ± 19 (6/6)* | *79 ± 1 (6/6)* | *371-591 (0/6)* |
| 10<sup>-4</sup> | *223 ± 15 (4/6)* | *361 ± 90 (4/6)* | *85 ± 1 (5/5)* | *399-518 (0/6)* |
| 10<sup>-5</sup> | *245 (1/6)* |  | *99 ± 2 (5/5)* | *367-636 (0/6)* |
| 10<sup>-6</sup> |  |  | *112 ± 2 (5/5)* |  |
| 10<sup>-7</sup> |  |  | *148 (1/5)* |  |

<sup>1</sup>10% brain material used as 10<sup>-0</sup> dilution.
<sup>2</sup>Days ± SE of the mean. For mice negative for PrP<sub>res</sub> accumulation, only the range of survival time is given.
<sup>3</sup>n/n<sub>0</sub>: Number of mice with neurological disease or positive for PrP<sub>res</sub> in the brain by immunoblotting/number of inoculated mice.
<sup>4</sup>tg110 line.
Data in italic are from [18].

Together, these observations indicate that PrP_Sc_ subassemblies segregation by SV-fractionation alters their replication efficiency in the PrP transmission barrier context, for reasons independent of their infectivity load. This suggests synergetic interactions between PrP_Sc_ subsets for an efficient heterospecies transmission.

### Dilution of LA21K *fast* prion-infected brain homogenate strengthens existing transmission barriers

LA21K *fast* prions are composed of at least two structurally PrP_Sc_ subsets in different proportion, each with distinct specific infectivity [23,15,18]. Dilution experiments constitute a relevant method to explore the contribution of each PrP_Sc_ subtype. Over a certain dilution factor, the minor population will be quasi-eliminated from the inoculum leading to explore the effect of the major species. The dilution approaches should also allow dissociating biochemical complex(es) between different PrP_Sc_ subsets due to equilibrium displacement toward dissociation [13].

We thus further analyzed the titration of LA21K *fast* prions in tgHa mice. Neat brain homogenate (20 μL 10%) containing LA21K *fast* prions induced disease in all tgHa mice with a mean ID of 153 days (Table 3; Additional file 1., supplementary Fig. 3a). At the 10_-1_ dilution, the mean ID increased to 181 days (6/6 mice affected). The limiting dilution value resulting in positive transmission established at the 10_-2_ dilution, with 2 out of 6 tgHa mice developing the disease at 192 and 230 days (Table 3). In homotypic transmission, LA21K *fast* limiting dilution value established at 10_-7_ (Table 3). There was thus a considerable 10_5_-fold reduction of this value in the heterotypic PrP transmission context. For comparison, in the absence of a transmission barrier, as for LA19K prions in tgBov mice, there was no significant variations in the limiting dilution value between the homotypic and heterotypic contexts (Table 3).

The inefficacy of LA21K *fast* diluted material to infect tgHa mice appeared further discrepant when considering the fold increase between the IDs at the lowest and at the limiting dilution. There was here a 1.37-fold increase for LA21K *fast* in tgHa mice (from 153 to 211 days, Table 3) whereas the mean±SD fold increase value observed in all the titrations we performed so far in our laboratory in homotypic conditions, including in tgHa mice, was statistically higher at 2.17±0.32 (p<0.05, One-sample z test; Fig. 4a). Applying this value to LA21K *fast* titration in tgHa mice would result in a theoretical limiting dilution value at ∼10_-5_ and a ∼330 days ID (Fig. 4b).

**Figure 4.**
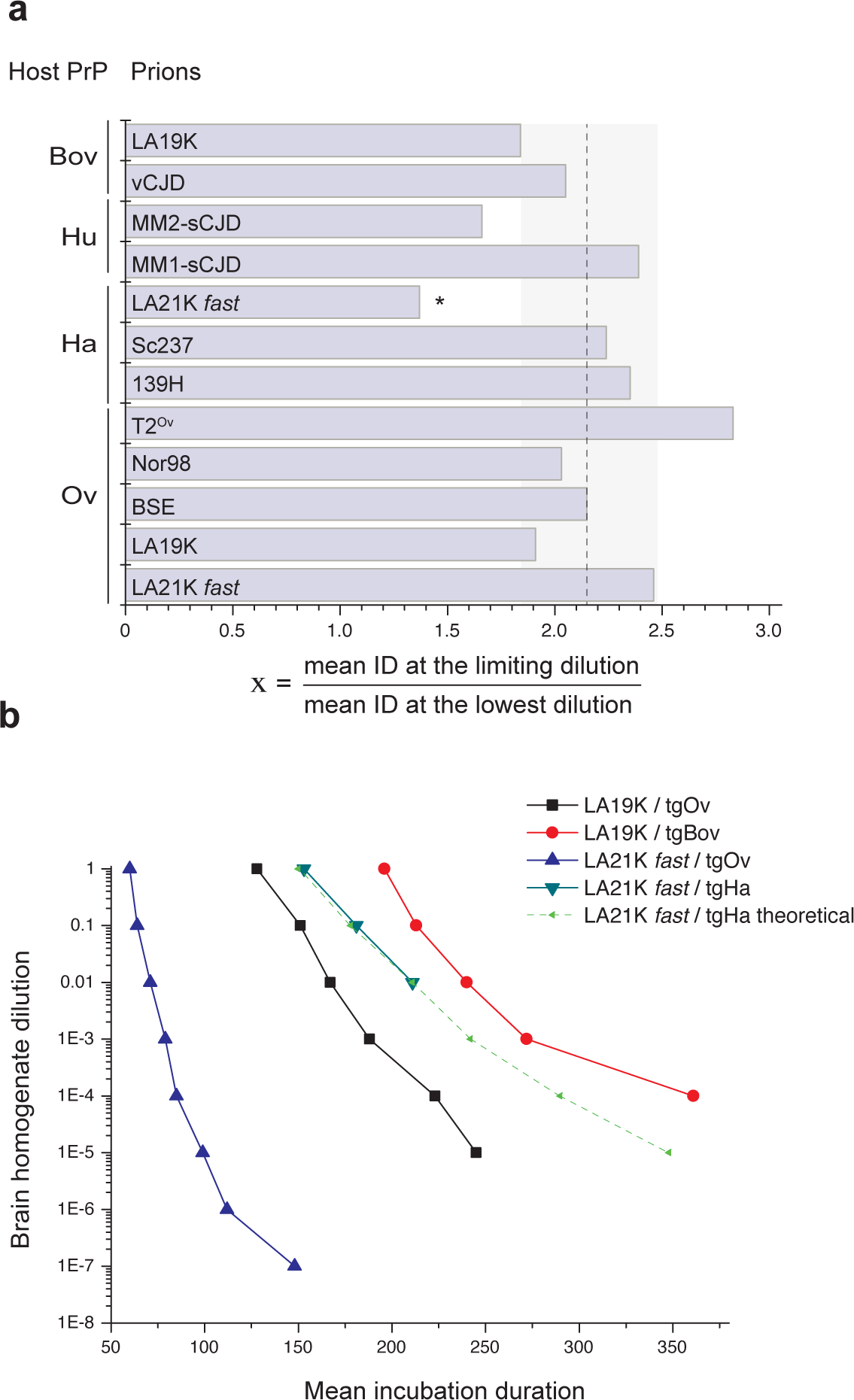
Aberrant titration of LA21K *fast* prion in hamster PrP mice as compared to other prion titrations in PrP transgenic mice. (**a**) Fold increase (x) between the mean IDs at the lowest and at the limiting dilution during prion titrations. The prions titrated and the reporter mice are indicated on the graph. The mean±SD value observed for all but the LA21K *fast*→tgHa titrations established at 2.17±0.32 (mean ± SD, dotted vertical line:mean; shadow square: SD). The 1.37-fold increase for LA21K *fast* in tgHa mice is significantly lower (*p<0.05, One-sample z test). (**b**) Titration curve of LA19K prions in tgOv and tgBov mice as compared to that of LA21K *fast* in tgOv and tgHa mice. The theoretical curve of LA21K *fast* prions in tgHa mice is the result of the 2.17-fold increase in the IDs at the lowest and at the limiting dilution.

Collectively, these data suggest that the dilution leads to the quasi elimination of at least one compound and/or the dissociation of a complex mandatory for the efficacy of the heterotypic transmission and consolidate our SV-based observations.

### PrP_C_ level and/or tissue environment imposes the species barrier stringency

When isolated L-BSE SV-fractions were transmitted to tgOv mice, only fractions 8 to 14 induced disease at complete or near-complete attack rate, with an overt delay compared to unfractionated material (Fig. 5a, Table 4). Mice inoculated with the top and bottom fractions transmitted the disease only erratically, suggesting unfavorable transmission conditions (Fig. 5a, Table 4). Isolated L-BSE assemblies had therefore altered transmission capacities in the heterotypic PrP transmission context, as observed with LA21K *fast* and 127S. The western blot profile on primary passage (Fig. 5b) and the strain phenotype obtained on 2_nd_ passage with PrP_res_-positive mouse brains indicated that C-BSE like prions has emerged from all positive transmissions (Additional File 1, supplementary Fig. 5; supplementary Table 1).

**Table 4.**
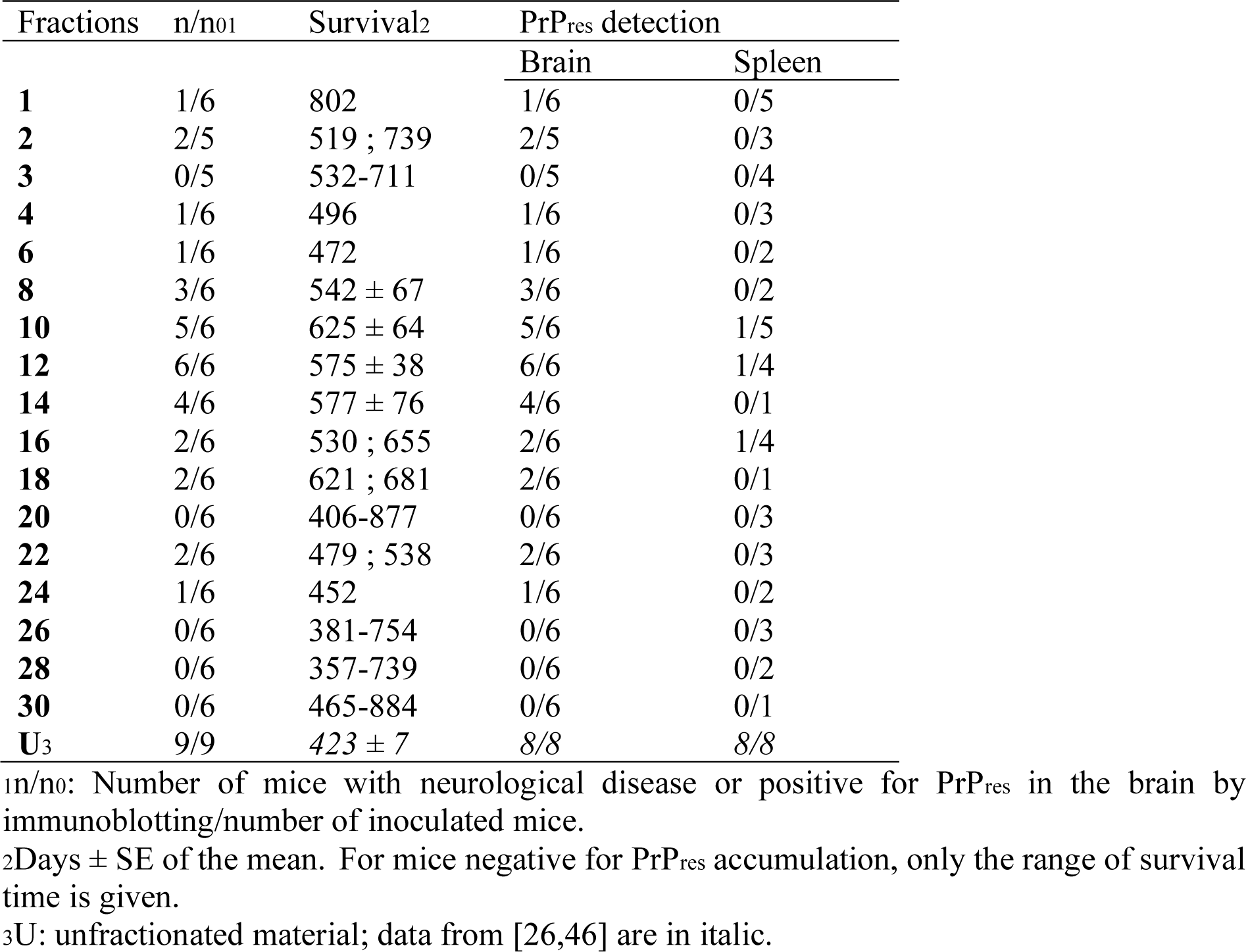
Survival time and PrP_res_ detection in the brain and spleen tissues of ovine PrP mice intracerebrally inoculated with L-BSE prions fractionated by sedimentation velocity.

| Fractions | n/n<sub>01</sub> | Survival<sup>2</sup> | PrP<sub>res</sub> detection |  |
| --- | --- | --- | --- | --- |
|  |  |  | Brain | Spleen |
| **1** | 1/6 | 802 | 1/6 | 0/5 |
| **2** | 2/5 | 519 ; 739 | 2/5 | 0/3 |
| **3** | 0/5 | 532-711 | 0/5 | 0/4 |
| **4** | 1/6 | 496 | 1/6 | 0/3 |
| **6** | 1/6 | 472 | 1/6 | 0/2 |
| **8** | 3/6 | 542 ± 67 | 3/6 | 0/2 |
| **10** | 5/6 | 625 ± 64 | 5/6 | 1/5 |
| **12** | 6/6 | 575 ± 38 | 6/6 | 1/4 |
| **14** | 4/6 | 577 ± 76 | 4/6 | 0/1 |
| **16** | 2/6 | 530 ; 655 | 2/6 | 1/4 |
| **18** | 2/6 | 621 ; 681 | 2/6 | 0/1 |
| **20** | 0/6 | 406-877 | 0/6 | 0/3 |
| **22** | 2/6 | 479 ; 538 | 2/6 | 0/3 |
| **24** | 1/6 | 452 | 1/6 | 0/2 |
| **26** | 0/6 | 381-754 | 0/6 | 0/3 |
| **28** | 0/6 | 357-739 | 0/6 | 0/2 |
| **30** | 0/6 | 465-884 | 0/6 | 0/1 |
| **U<sub>3</sub>** | 9/9 | 423 ± 7 | 8/8 | 8/8 |

<sup>1</sup>n/n<sub>0</sub>: Number of mice with neurological disease or positive for PrP<sub>res</sub> in the brain by immunoblotting/number of inoculated mice.
<sup>2</sup>Days ± SE of the mean. For mice negative for PrP<sub>res</sub> accumulation, only the range of survival time is given.
<sup>3</sup>U: unfractionated material; data from [26,46] are in italic.

**Figure 5.**
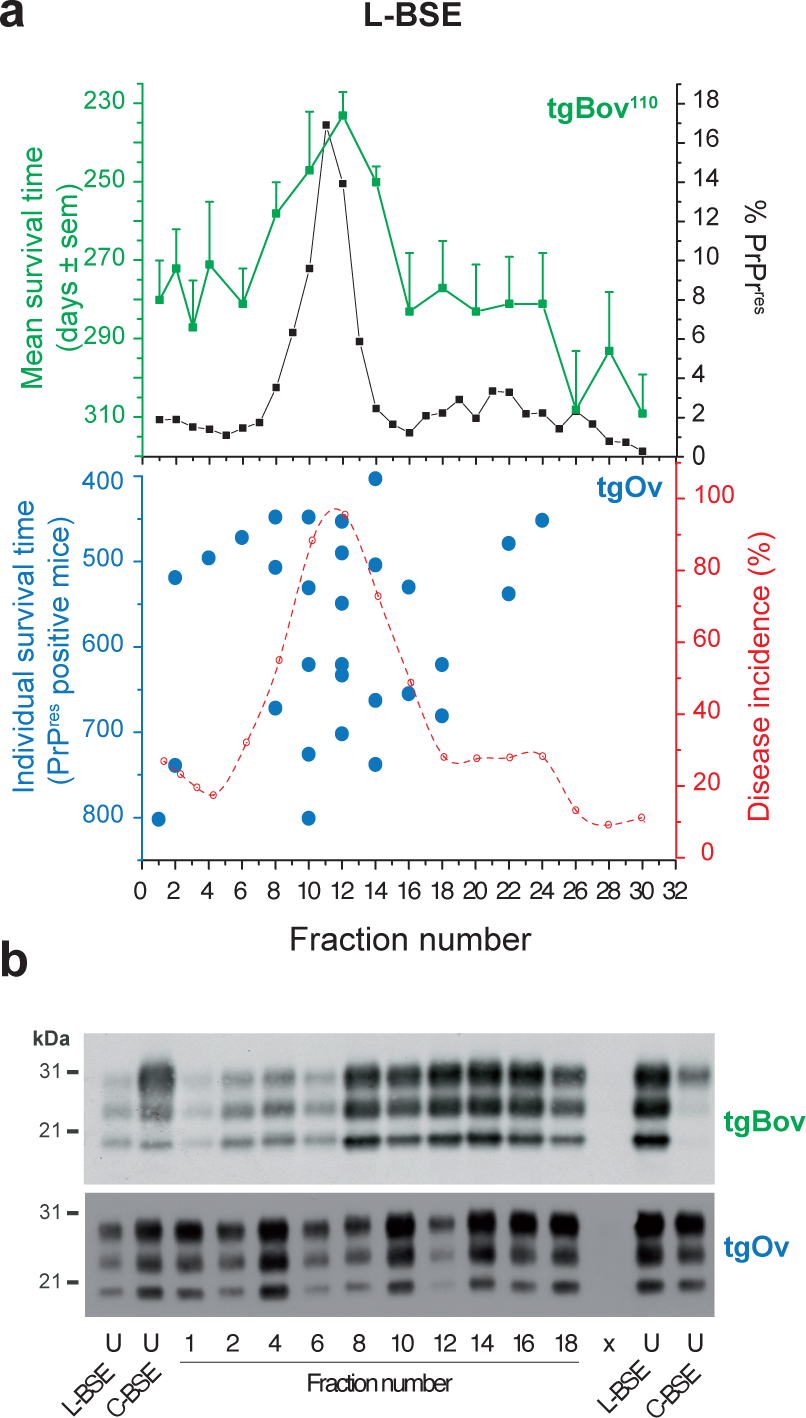
Capacity of size-fractionated PrP_Sc_ assemblies to propagate onto a new host PrP sequence in the presence of an ‘intermediate’ transmission barrier and with mutation. (**a**) SV profiles of PrP_res_ (black line) and infectivity in the homotypic PrP (green line) and heterotypic PrP (blue dots) transmission context. Brain homogenates from cattle infected with L-BSE prions were solubilized before fractionation by SV. The collected fractions were analyzed for PrP_res_ content by immunoblot and for infectivity by an incubation time bioassay in tgBov and tgOv mice. The disease incidence in tgOv mice is presented on the right red graph. Because of the reduced penetrance of the disease in tgOv mice, the mean individual ID of the PrP_res_ positive mice are shown as blue dots. (**b**) PrP_res_ electrophoretic profiles in the brains of tgOv and tgBov mice inoculated with size-fractionated L-BSE PrP_Sc_ assemblies. The profiles obtained with unfractionated (U) material from L-BSE or classical BSE (C-BSE) are shown for comparison.

Previously, we reported that lymphotropic prions replicated easier in the spleen than in the brain in both homotypic and heterotypic PrP context, despite 20-fold lowered PrP_C_ levels [27,45]. This makes bioassays based on prion detection in the mouse spleen a highly sensitive method to detect low dose of lymphotropic prions [45]. We thus examined further the capacity of SV-fractionated L-BSE prions to propagate in the heterotypic PrP context by examining tgOv spleens for PrP_res_ content after inoculation with the different fractions. Of the 49 tgOv spleens analyzed, only 3 (6%) accumulated low levels of PrP_res_ (Table 4). Of the 13 mice for which both the brains and spleens were analyzed for PrP_res_ content and the brain was positive (notably fractions 10-12 with 100% attack rate), only 3 had also positive spleens (23%). For comparison, all the spleens analyzed after inoculation of unfractionated L-type prions were strongly PrP_res_-positive from the first passage onward (Table 4, [46]).

Strikingly, prion replication in the spleen was still altered on secondary passage of SV-fractionated L-BSE assemblies. While all the spleens analyzed (n=15) were PrP_res_-positive, the levels of accumulation were still reduced by ∼4-fold as compared to those observed on serial transmission of unfractionated material (Fig. 6a-b). It may be noted that in homotypic transmissions, fractionating brain material had no significant influence on spleen colonization by prions, with regards to the number of positive spleens and PrP_res_ accumulation levels (Fig. 6b), indicating that, alone, the size of the infectious particles or PrP_Sc_ subpopulation segregation were not causal. Therefore, we can conclude from the impairment observed over two serial passages that the low barrier penetrance of the isolated fractions is not due to a low infectivity load and that size fractionation impairs L-BSE replication in the spleen to an extent that is higher than in the brain.

**Figure 6.**
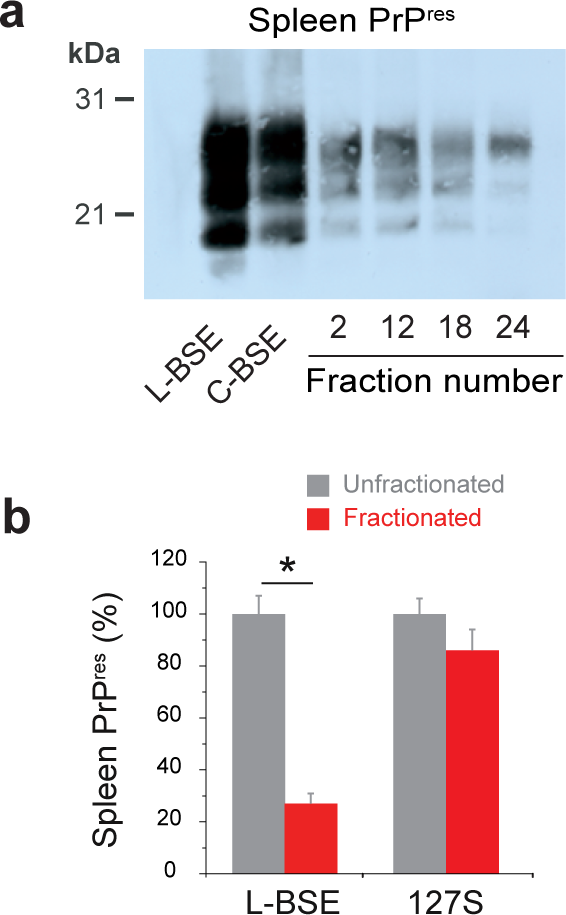
Capacity of size-fractionated L-BSE prions to colonize the spleens of mice expressing homotypic or heterotypic PrP_C_. (**a**) PrP_res_ detection in the spleens of tgOv mice inoculated with unfractionated L-BSE or C-BSE prions (primary passage) or with fractionated L-BSE prions (second passage, fractions used for passaging are indicated). (**b**) PrP_res_ accumulation levels in the spleens of tgOv mice inoculated with unfractionated (2_nd_ passage) or SV-fractionated L-BSE ((all fractions combined), 2_nd_ passage) prions as compared to unfractionated or SV-fractionated 127S prions. (\**p*<0.05, Mann-Whitney test).

## Discussion

As conventional pathogens, prions are subject to adaptation and evolution. The concept of molecular quasispecies, as defined by Eigen in 1979 [47], has been applied to prions [5,48] to reconcile the structural diversity of prion assemblies (i.e. prion structural landscape) to the preferential selection based on the ‘best replicator’ selection concept of certain subassemblies during prion adaptation to new host or to a new environment/replicative conditions [49,27]. To refine this ‘best replicator’ hypothesis [50], we tested the capacity of isolated PrP_Sc_ subassemblies within a given prion strain to adapt to a new host. We show here with multiple PrP transgenic mouse models inoculated with both field-derived and biologically cloned prions, that cross-species prion transmission does not strictly involve the selection of an existing subpopulation of optimized PrP_Sc_ assemblies. Rather, our observations suggest that the key determinant is a synergy between structurally distinct PrP_Sc_ subassemblies.

There was a positive correlation between the magnitude of the transmission barrier and the difficulty for SV-individualized PrP_Sc_ assemblies to transmit the disease in the heterotypic PrP conditions. L-BSE prions fractionation significantly delayed priogenesis in the brain and even more negatively in the spleen of the challenged tgOv mice. Fractionated LA21K *fast* scrapie prions replicated asymptomatically in a very small proportion of tgHa mice and fractionated 127S prions failed to propagate in these mice. As shown previously and also demonstrated here, LA21K *fast* and 127S scrapie prion strains are composed of at least two structurally distinct PrP_Sc_ subpopulations according to their specific infectivity values (Fig. 3a and [15,18,23]). The loss of transmissibility of separately taken PrP_Sc_ assemblies to tgHa mice cannot be attributed to a global decrease of infectivity levels due to the solubilization/fractionation method as we previously reported that the cumulated infectivity of the SV fractions did not differ significantly from that present in the loaded material [18]. Further, for LA21K *fast*, the relative infectivity titer of the top ‘most infectious’ fractions was sufficiently high to induce disease in tgHa. The fact that PrP_Sc_ assemblies segregation affects prion heterospecies transmission suggests a synergy between PrP_Sc_ subassemblies to overcome the species barrier. This hypothesis implicates an unprecedented notion of complementation between structurally distinct prion assemblies.

While the molecular mechanisms of such synergy between different PrP_Sc_ subsets remain to be elucidated, some simple biochemical considerations can bring basements on the biochemistry of the process. It implicates interactions between structurally distinct PrP_Sc_ subsets in order to create a new structural information absent in each individual PrP_Sc_ population and leading to heterologous PrP_C_ integration. During the early stage of prion replication, we showed that two sets of structurally distinct PrP_Sc_ assemblies are generated, in fractions 1-5 and 10-18 [20]. These sets are defined by their elementary subunit [13] as suPrP_A_ and suPrP_B_, respectively [27]. During the early step of prion replication, suPrP_A_ and suPrP_B_ form a suPrP_A_:suPrP_B_ heterocomplex involved in a secondary autocatalytic templating process generating *de novo* the formation of suPrP_B_, in a PrP_C_-dependent manner [27]. The synergetic effect between different PrP_Sc_ subsets can reside in the formation of this complex, making therefore this secondary templating process a pivot of prion adaptation to a new host. In the heterotypic PrP transmission context, the suPrP_A_:suPrP_B_ complex (likely present in the inoculum) may incorporate heterologous PrP_C_, leading to the formation of a *de novo* suPrP_B_* formed by the heterologous PrP_C_ with a new templating interface different from the inoculum suPrP_B_ (Fig. 7). This first event would be a limiting step of the adaptation process. The entrance of suPrP_B_* in the autocatalytic cycle makes its formation highly cooperative. Based on these assumptions, if the stability of the complex is high enough (i.e., low dissociation constant), the reconstitution of the initial composition by mixing different PrP_Sc_ subsets should allow recovering the transmission efficiency. This approach has been unsuccessfully attempted in the case of LA21K *fast*. This fail could be due to the weakness of the suPrP_A_:suPrP_B_ complex, as previously observed [20]. Indeed, size exclusion chromatography analysis of the 127S suPrP_A_:suPrP_B_ complex revealed that this last is highly labile and can be observed only in conditions where the concentration of suPrPA and suPrPB were high.

**Figure 7.**
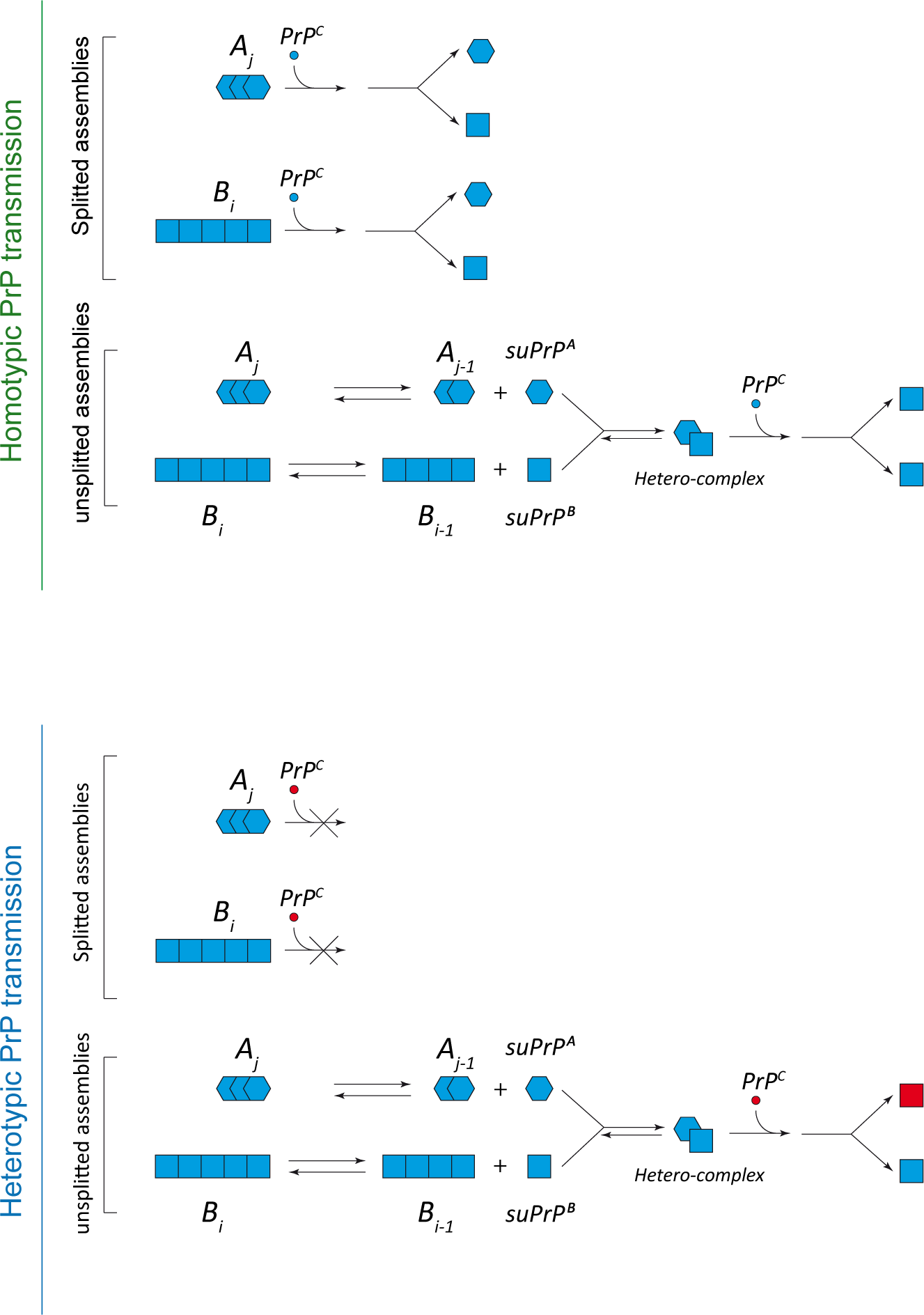
The formation of heterocomplex between structurally distinct PrP_Sc_ subsets may drive the conversion of heterologous PrP_C_ by a secondary templating pathway process. (**a**) In the homotypic PrP transmission context, prion replication involves two templating pathways: a primary templating pathway where structurally distinct PrP_Sc_ subsets, taken individually (A_j_ and B_i_), are able to perpetuate the strain information and a PrP_C_-dependent autocatalytic secondary templating pathway contributing to structural diversification, which requires the formation of a heterocomplex between the elementary subunits of two structurally distinct sets of PrP_Sc_ assemblies (suPrP_A_ and suPrP_B_) [20]. (**b**) This autocatalytic secondary templating pathway could drive the incorporation of heterologous PrP_C_ during the species barrier passage, leading to *de novo* generation of suPrP_B_ (in red) more prone to replicate/propagate in the new host. PrP_Sc_ assemblies segregation or dilution could drastically affect the formation of the suPrP_A_:suPrP_B_ complex, compromising this secondary templating pathway.

The existence of a complex such as suPrP_A_:suPrP_B_ within the inoculum is further supported by the striking observation that endpoint dilutions aberrantly impaired LA21K *fast* prions transfer to tgHa mice (1000-fold decreased efficacy); the dilution steps would make certain PrP_Sc_ subpopulations disappearing given their ratio/amount and/or induce the dissociation of an existing complex [51,15]. While cross-species transmission of prions has rarely been done at high dilution, it may be noted that comparing the limiting dilution values of MM1 sporadic CJD prions (homozygous for Met at codon 129) in transgenic mice expressing human PrP with either Met or Val at codon 129 resulted in 103-fold decrease [52], a negative impact quantitatively comparable to our observations.

If the synergy between structurally different PrP_Sc_ subsets involves the formation of a suPrP_A_:suPrP_B_ heterocomplex and the PrP_C_-dependent secondary templating pathway, the magnitude of the species barrier would thus depend both on the strength of the heterocomplex and on the amount of heterotypic PrP_C_ which defines the rate of the secondary templating and its cooperativity aspect [20]. Such prominent role of PrP_C_ is experimentally supported by the observation that L-BSE prion adaptation is more impaired in tg338 mouse spleen than brain, despite a potentially favored local environment for heterotypic conversion as prion species barrier is lower in the spleen than in the brain [27]. The lower concentrations of PrP_C_ [53] in the spleen [27] and/or the different structure (including the post-translational modifications) may impact the secondary templating pathway and its autocatalytic nature.

## Conclusion

Together, our data would expand the prion quasispecies concept to an ensemble of macromolecular assemblies, -not necessarily associated to different substrains-, that complement each other to adapt tissue-specific selection pressure and extend prion host range. Such ‘epistatic’ behavior may provide new fundamental principles for understanding prion distinctive properties to transfer between species [54] and can explain the relative inability of recombinant PrP to generate high-titer prions [55], unless submitted to intense mechanisms of polymerization/fragmentation which may generate aggregate polydispersity and/or a larger portfolio of conformations [56] for complementation.

## List of Abbreviations

Bov: bovine
BSE: bovine spongiform encephalopathy
CJD: Creutzfeldt-Jakob disease
Ha: hamster
Ov: ovine
PrP: prion protein
PK: proteinase K
PrP_res_: proteinase K resistant PrP_Sc_
PrP_Sc_: pathological/abnormal form of the prion protein
suPrP: PrP elementary brick
SV: sedimentation velocity
tg: transgenic

## Declarations

### Ethics approval

All the experiments involving animals were carried out in strict accordance with EU directive 2010/63 and were approved by INRA local ethics committee (Comethea; permit numbers 12-034 and 15-056).

### Consent for publication

Not applicable.

### Availability of data and material

All data supporting our findings are presented in the main paper and additional files.

## Acknowledgments

We thank the staff of Infectiology of fishes and rodent facilities (INRA, Jouy-en-Josas, France; doi:10.15454/1.5572427140471238E12) for animal care.

## Author’s contribution

Conceptualization, AIE, FL, PT, MM, HL, HR, and VB; Data curation and Analysis, AIE, FL, PT, MM, LH, FR, JMT, HL, HR, and VB; Resources, HL, JMT, HR and VB; Writing – Original Draft, AIE, HR, and VB; Writing – Review and Editing, AIE, HR, and VB; Visualization, AIE, HR and VB; Supervision, VB; Funding Acquisition, HR and VB; All authors read and approved the final manuscript.

