## Supplemental data for "Complementation between pathological prion protein subassemblies to cross existing species barriers"

**Additional File 1**

As transgenic model of prion transmission without species barrier, we passed LA19K scrapie prions from ovine PrP mice to bovine PrP mice (tg540 [1] or tg110 [2] line; tgBov). Iterative transmission and retrotransmission of LA19K prions occurred in these lines without drastic evolution of time to disease onset, PrP$^{Sc}$ biochemical properties and PrP$^{Sc}$ distribution in brain and extraneural tissue (additional File 1, supplementary Fig. 1). Therefore, the magnitude of the transmission barrier was ~1 (Fig. 1a), suggesting that LA19K strain structural determinant belongs to the structural landscape of bovine PrP$^{C}$.

As model of prion transmission with species barrier and 'mutation', we passed L-BSE prions onto tgOv mice (additional File 1, supplementary Fig. 2 and [3]). L-BSE prions adaptation to tgOv mice required iterative passages, resulting in a transmission barrier magnitude value of ~3 (Fig. 1a). This adaptation was accompanied by a "mutation" of L-BSE prions into classical-like BSE (C-BSE) prions [3]. However, L-BSE prions were readily reisolated on back-passage to tgBov mice, without any significant transmission barrier, as based on the mean ID and regain of L-type phenotypic identity (Fig. 1a, additional File 1, supplementary Fig. 2a-c). The transmission barrier was thus qualified as 'medium'.

As models of prion transmission with stronger species barrier, we passed either LA21K *fast* or 127S scrapie prions onto hamster PrP mice (tg7 line [4]; tgHa). These two prion strains can adapt to tgHa mice and be reisolated without strain identity change in tgOv mice (additional File 1, supplementary Fig. 3-4). Yet, at each of these crossings, iterative passages were necessary to attain minimal IDs (additional File 1, supplementary Fig. 3), resulting in transmission barrier magnitudes of ~3.5 (tgOv $\rightarrow$ tgHa) and ~6-fold (tgHa $\rightarrow$ tgOv) (Fig. 1a).

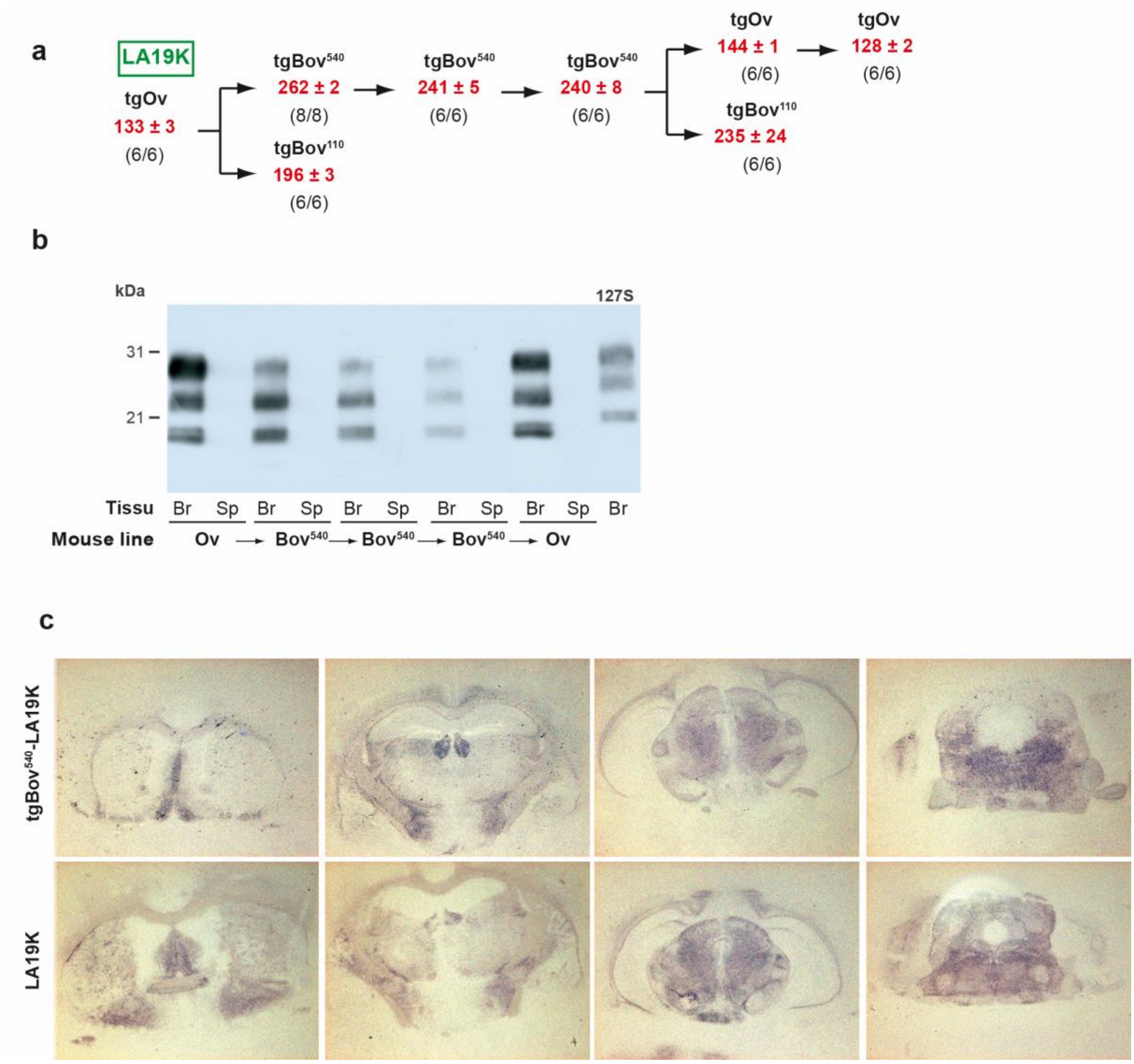


**Supplementary Figure 1. Propagation of LA19K scrapie prions to bovine PrP mice without apparent transmission barrier**

(**a**) Serial propagation (intracerebral route) of LA19K scrapie prions [5,6] from ovine PrP (tgOv; tg338 line) to bovine PrP (tgBov; tg540 [1] (Bov$_{540}$) or tg110 [2] (Bov$_{110}$) lines) transgenic mice and retrotransmission to ovine PrP mice. At each passage, the mean ± SEM incubation durations (ID; in red) and the attack rate are indicated. On primary passage to tgBov$_{540}$ mice, the mean ID established at approximately 260 days and decreased modestly to 240 days on further passaging. tgBov$_{540}$-derived LA19K prions were as virulent as LA19K prions on retrotransmission to tgOv mice; disease occurred at full attack rate in 144 days and in 128 days on second passage.

(**b**) PK-resistant $PrP_{Sc}$ ($PrP_{res}$) detection and electrophoretic banding pattern on serial passage of LA19K prions to tgBov mice (tg540 line) and on backpassage to tgOv mice. Western blot analyses were made in brain (Br) and spleen (Sp) tissues of the diseased mice. Scrapie 127S $PrP_{res}$ electrophoretic mobility in tgOv mouse brain is shown for comparison (21 kDa pattern [7] as referred to the unglycosylated band of $PrP_{res}$). In tgOv as in tgBov mice, LA19K scrapie prion exhibited a 19kDa electrophoretic signature with regard to unglycosylated $PrP_{res}$, and $PrP_{res}$ was not detected in the spleen. On back passage to tgOv mice, these characteristics were maintained.

(**c**) Neuroanatomical pattern of $PrP_{res}$ deposition in tgOv mice inoculated with either LA19K or $tgBov_{540}$-derived LA19K prions. Representative histoblots of antero-posterior coronal brain sections (12F10 antibody) at the level of the septum, hippocampus, midbrain and brainstem (from left to right) are shown.

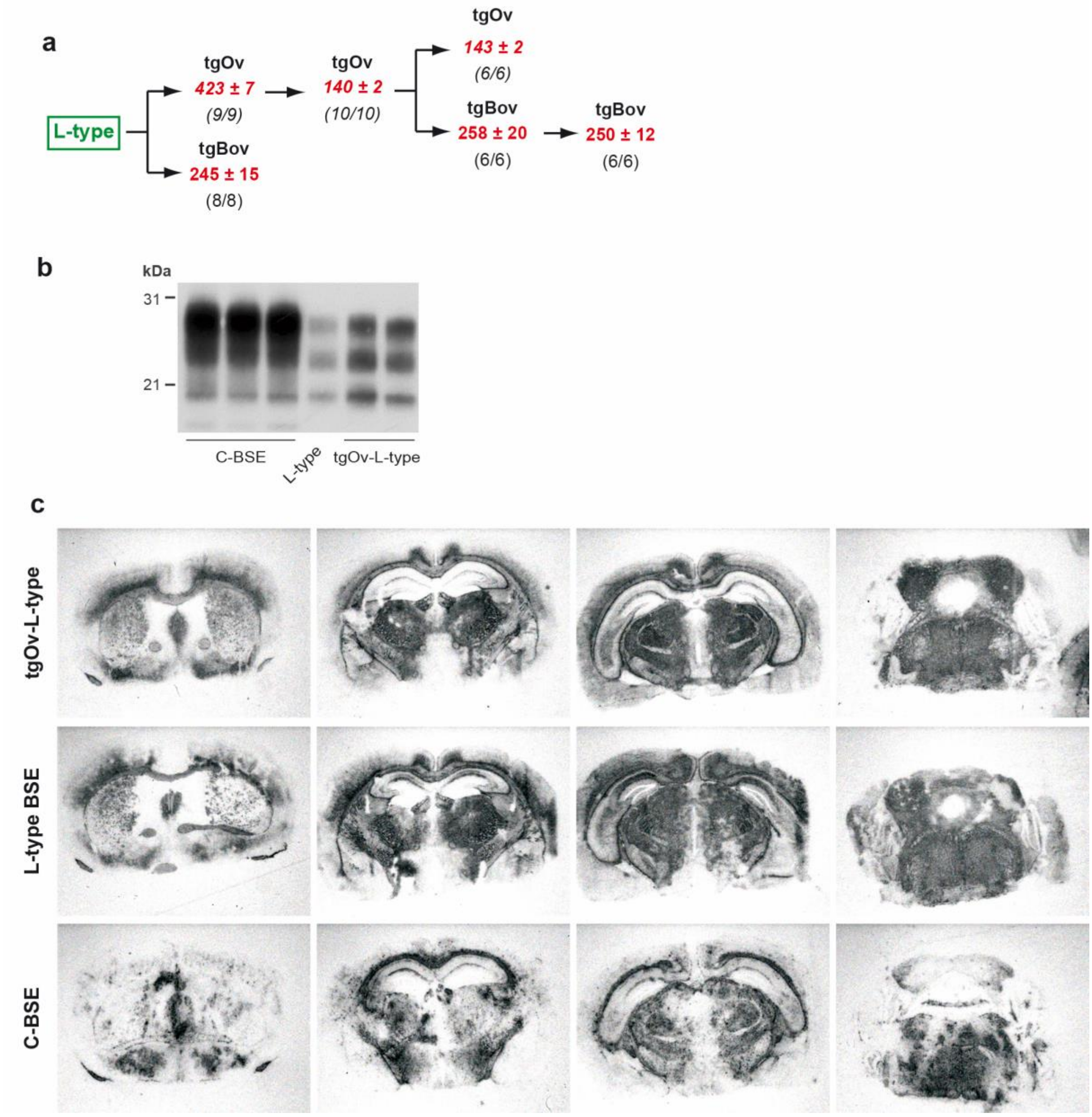


**Supplementary Figure 2. Reisolation of L-type BSE on back passage from tgOv to tgBov mice**

Serial propagation by intracerebral route of L-type BSE [3,8] from cattle to tgOv mice and reisolation in bovine PrP mice (tg110 line). At each passage, the mean ± SEM ID (in red) and the attack rate are indicated. On primary transmission to tgOv mice, all the inoculated mice developed the disease with a mean ID of approx. 420 days. On further passaging, the ID was reduced by 3-fold, establishing at 140 days (data published in reference [3]). tgOv-derived L-BSE prions were as rapidly pathogenic as cattle L-type prions on back-passage to tgBov mice, suggesting absence of transmission barrier on return to the parental $PrP_C$ sequence (Fig. 1a).

(**b**) $PrP_{res}$ banding pattern in the brains of tgBov mice inoculated with tgOv-derived L-BSE prions. The profile of classical BSE (C-BSE) and of L-BSE in tgBov is shown for comparison. Note that tgOv-derived L-BSE prions reacquired a L-BSE $PrP_{res}$ phenotype on back-passage to tgBov mice.

(**c**) Comparison of $PrP_{res}$ deposition in the brains of tgBov mice inoculated with tgOv-derived L-BSE prions, L-type BSE and C-BSE. Representative histoblots of antero-posterior coronal brain sections ($2_{nd}$ passage, 12F10 antibody) at the level of the septum, hippocampus, midbrain and brainstem (from left to right) are shown. Note that tgOv-derived L-BSE $PrP_{res}$ distribution resembled that of L-BSE in tgBov mice.

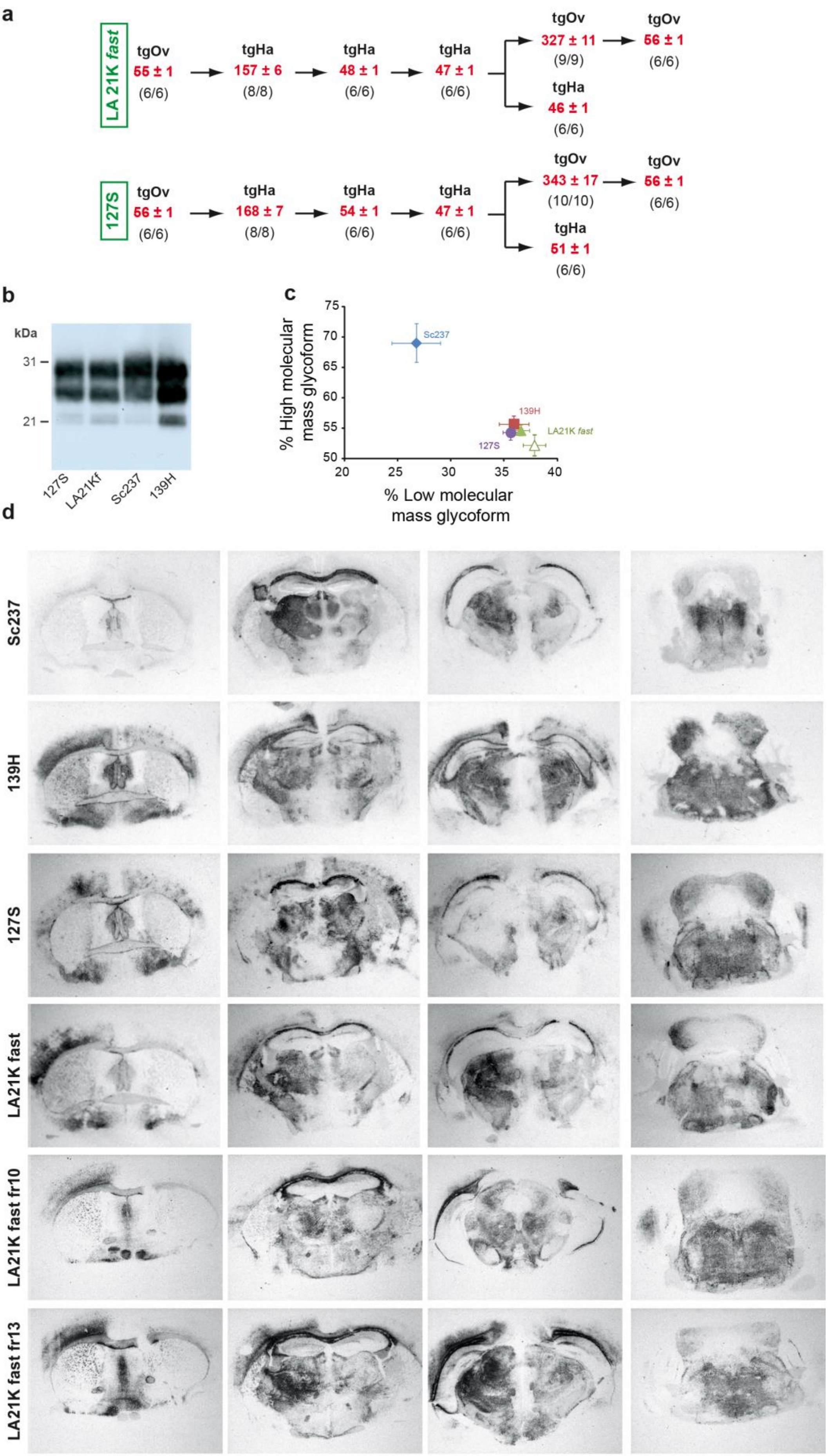
a
LA 21K fast
tgOv
55 ± 1
(6/6)
tgHa
157 ± 6
(8/8)
tgHa
48 ± 1
(6/6)
tgHa
47 ± 1
(6/6)
tgOv
327 ± 11
(9/9)
tgOv
56 ± 1
(6/6)
tgHa
46 ± 1
(6/6)
127S
tgOv
56 ± 1
(6/6)
tgHa
168 ± 7
(8/8)
tgHa
54 ± 1
(6/6)
tgHa
47 ± 1
(6/6)
tgOv
343 ± 17
(10/10)
tgOv
56 ± 1
(6/6)
tgHa
51 ± 1
(6/6)
b
kDa
31
21
127S
LA21Kf
Sc237
139H
c
% High molecular mass glycoform
% Low molecular mass glycoform
Sc237
139H
LA21K fast
127S
d
Sc237
139H
127S
LA21K fast
LA21K fast fr10
LA21K fast fr13

**Supplementary Figure 3. Propagation of LA21K *fast* and 127S scrapie prions in hamster PrP mice points to a substantial transmission barrier**

(**a**) Serial propagation (intracerebral route) of LA21K *fast* and 127S scrapie prions [5,7,6] from ovine PrP (tgOv; tg338 line) to hamster PrP (tgHa; tg7 line[4]) transgenic mice and retrotransmission to ovine PrP mice. At each passage, the mean ± SEM ID (in red) and the attack rate are indicated. On primary passage of LA21K *fast* and 127S to tgHa mice, all the inoculated mice developed the disease with mean IDs of 157 and 168 days, respectively. On further passaging, the ID was reduced by 3-fold, establishing at 46 and 51 days, respectively. The mean IDs of tgHa-derived LA21K *fast* and 127S were reminiscent of Sc237 prions in tgHa mice (45 ± 1 days [6]). tgHa-derived LA21K *fast* and 127S prions were pathogenic on back-passage to tgOv mice. It took however two passages to restore a mean ID superimposable to the parental prions (see supplementary Fig. 4 for characterization of strain phenotype). Such magnitude in the reduction of the mean ID on adaptation to the hamster PrP sequence and on back passage (Fig. 1a) points to a substantial transmission barrier [9].

(**b**) $PrP_{res}$ electrophoretic banding pattern and (**c**) mean (± SEM) ratios of high- to low-molecular mass $PrP_{res}$ glycoforms (*n=3* brains analyzed in triplicate at the $3_{rd}$ passage, (**c**)) in the brain of tgHa mice challenged with unfractionated (plain triangle) or fractionated (open triangle) LA21K *fast* prions and with unfractionated 127S prions (plain circle). Sc237 (plain diamond) and 139H (plain square) hamster prion strains are shown for comparison. $PrP_{res}$ glycoform ratio of LA21K *fast* and 127S in tgHa mice resembled that of 139H $PrP_{res}$ in tgHa mice.

(**d**) Neuroanatomical pattern of $PrP_{res}$ deposition in tgHa mice inoculated with LA21K *fast* (unfractionated or after fractionation, $2_{nd}$ passage) and unfractionated 127S prions. Representative histoblots of antero-posterior coronal brain sections ($4_{th}$ passage, 3F4 antibody) at the level of the septum, hippocampus, midbrain and brainstem (from left to right) are shown. The pattern observed with LA21K *fast* and 127S resembled that of 139H, except the more pronounced presence of $PrP_{res}$ deposits in the corpus callosum which recalls Sc237 prions. The deposition pattern observed with unfractionated and fractionated LA21K *fast* material were superimposable.

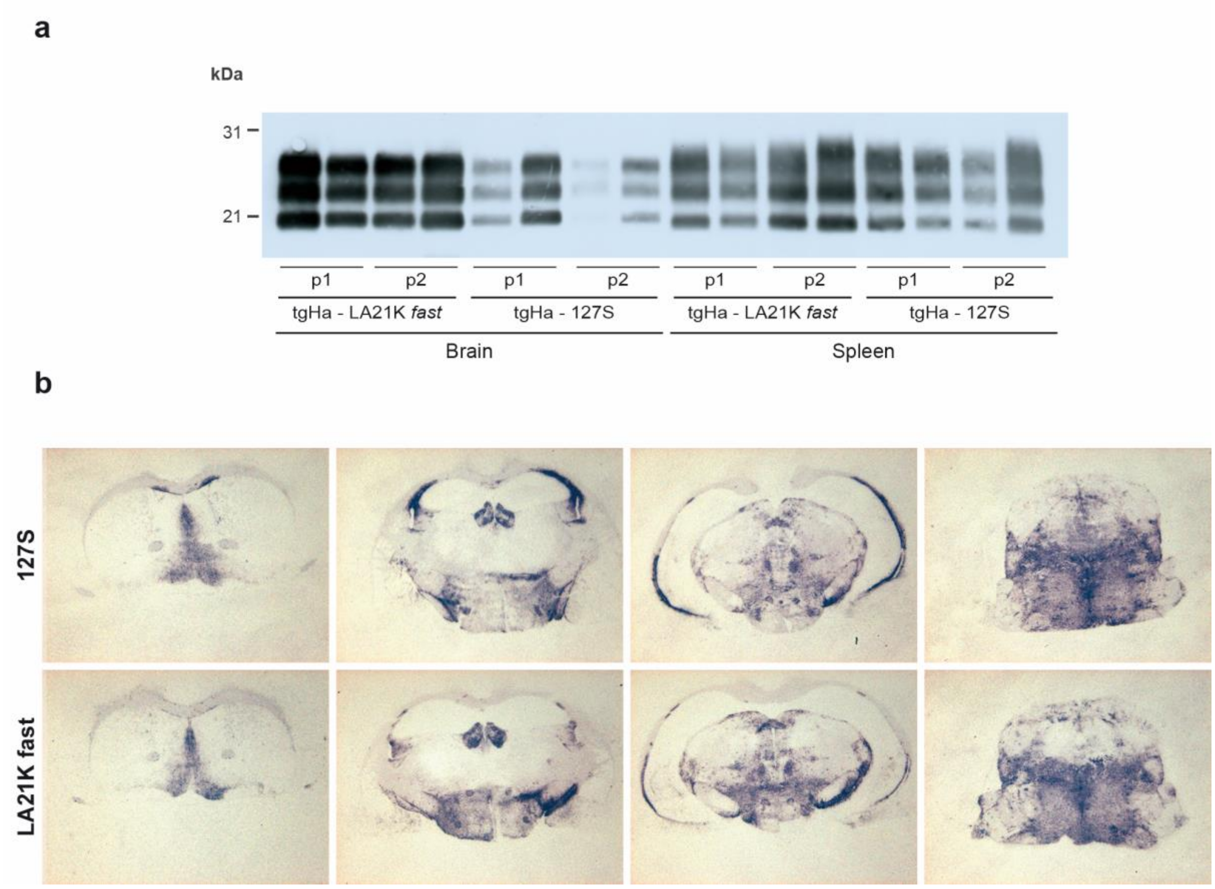


**Supplementary Figure 4. Reisolation of LA21K *fast* and 127S strains on back passage from tgHa to tgOv mice**

LA21K *fast* and 127S prion serially propagated onto tgHa mice (4th passage, supplementary Fig. 3a) were transmitted back to tgOv mice (intracerebral route).

(**a**) $PrP_{res}$ detection and banding pattern of tgHa-derived LA21K *fast* and 127S prions on two back-passages (p1, p2) in tgOv mice. Western blot analysis was made in brain and spleen tissues of the diseased mice. The profiles obtained were typical of the parental strains [7,6].

(**b**) Neuroanatomical pattern of $PrP_{res}$ deposition in tgOv mice inoculated with tgHa-derived LA21K *fast* and 127S prions. Representative histoblots of antero-posterior coronal brain sections (2nd passage, 12F10 antibody) at the level of the septum, hippocampus, midbrain and brainstem (from left to right) are shown. The profiles observed were typical of the parental strains [7,6].

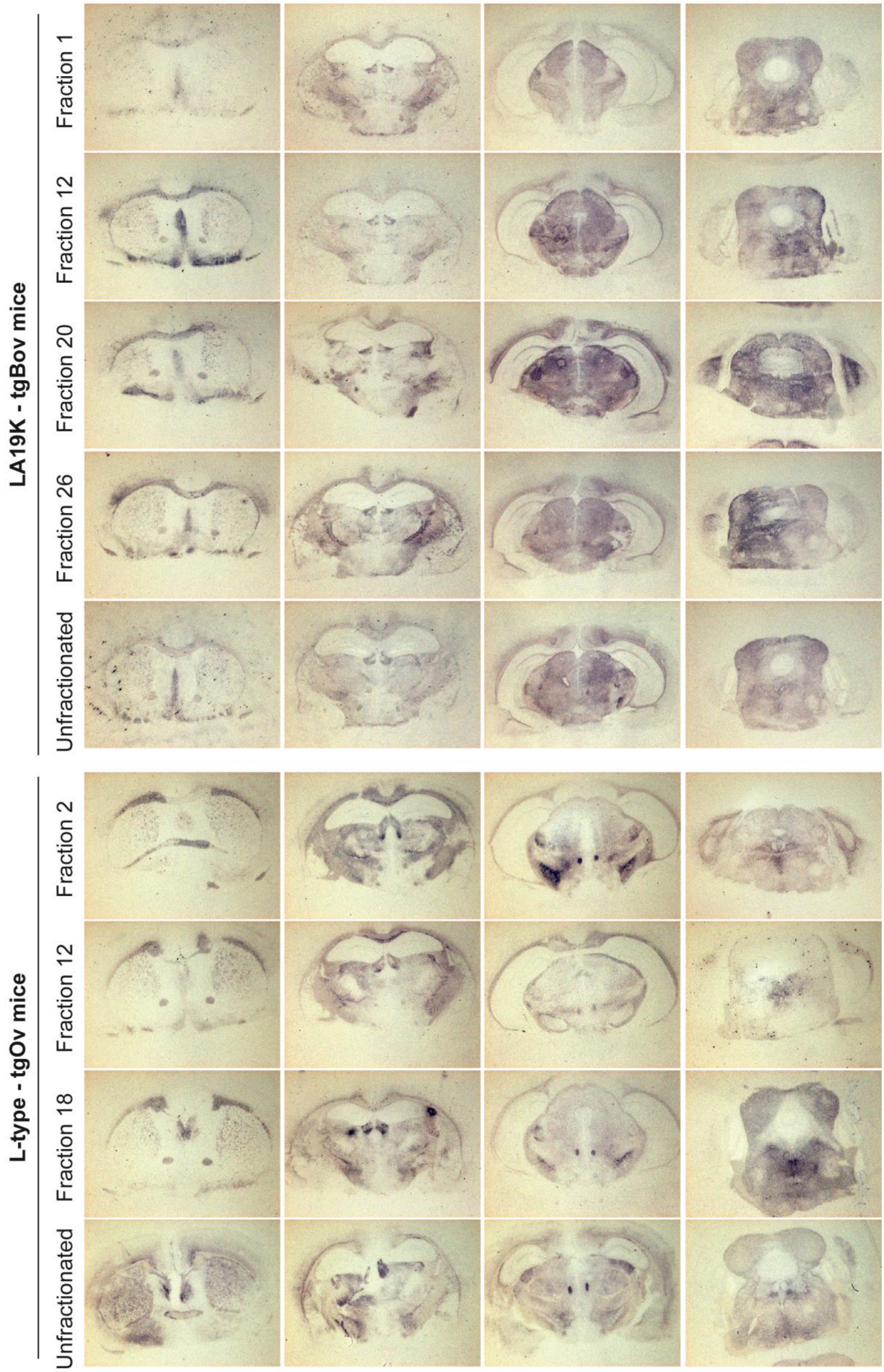
LA19K - tgBov mice
Fraction 1
Fraction 12
Fraction 20
Fraction 26
Unfractionated
L-type - tgOv mice
Fraction 2
Fraction 12
Fraction 18
Unfractionated

**Supplementary Figure 5. Neuroanatomical pattern of $PrP_{res}$ deposition in tgBov and tgOv mice inoculated with fractionated or unfractionated LA19K and L-BSE prions**

tgBov (tg110 line) and tgOv were inoculated with fractionated or unfractionated LA19K and L-type BSE prions, respectively. Representative histoblots of antero-posterior coronal brain sections (1st passage for LA19K, 2nd passage for L-BSE, 12F10 antibody) at the level of the septum, hippocampus, midbrain and brainstem (from left to right) are shown.

**Supplementary Table 1. Second passage by intracerebral route of SV fractionated L-BSE prions in ovine PrP mice**

| **Fractions** | L-BSE | |
|---|---|---|
| (mouse no.[1]) | $n/n_0$[2] | Survival[3] |
| **2** (no. 2) | 7/7 | 162 ± 3 |
| **12** (no.3) | 6/6 | 157 ± 3 |
| **18** (no. 3) | 6/6 | 151 ± 1 |
| **24** (no.4) | 6/6 | 193 ± 6 |

[1]mouse number inoculated.

[2]$n/n_0$: Number of mice with neurological disease and positive for $PrP_{res}$ in the brain by immunoblotting/number of inoculated mice (the number of the mouse inoculated is indicated).

[3]Mean survival time (days ± SE of the mean).
